# DectiSomes: Glycan Targeting of Liposomal Drugs Improves the Treatment of Disseminated Candidiasis

**DOI:** 10.1101/2021.07.27.454088

**Authors:** Suresh Ambati, Tuyetnhu Phan, Zachary A. Lewis, Xiaorong Lin, Richard B. Meagher

## Abstract

*Candida albicans* causes life-threatening disseminated candidiasis. Individuals at greatest risk have weakened immune systems. An outer cell wall, exopolysaccharide matrix, and biofilm rich in oligoglucans and oligomannans help *Candida spp.* evade host defenses. Even after antifungal drug treatment the one-year mortality rate exceeds 25%. Undoubtedly there is room to improve antifungal drug performance. The mammalian C-type lectin pathogen receptors Dectin-1 and Dectin-2 bind to fungal oligoglucans and oligomannans, respectively. We previously coated amphotericin B-loaded liposomes, AmB-LLs, pegylated analogs of AmBisome, with the ligand binding domains of these two Dectins. DectiSomes, DEC1-AmB-LLs and DEC2-AmB-LLs, showed two distinct patterns of binding to the exopolysaccharide matrix surrounding *C. albicans* hyphae grown in vitro, while untargeted AmB-LLs did not bind. DectiSomes were preferentially associated with fungal colonies in the kidneys. In a neutropenic mouse model of candidiasis, DEC1-AmB-LLs and DEC2-AmB-LLs delivering only one dose of 0.2 mg/kg AmB significantly reduced the kidney fungal burden several fold relative to AmB-LLs, based on either colony forming units (P= 0.013 to 8.8 × 10^-5^) or quantitative PCR of fungal rRNA ITS (P= 5.5×10^-5^ to 3.0×10^-10^). DEC1-AmB-LLs and DEC2-AmB-LLs significantly increased the percent of surviving mice relative to AmB-LLs. Dectin-2 targeted anidulafungin loaded liposomes and AmBisomes, DEC2-AFG-LLs and DEC2-AmBisome reduced fungal burden in the kidneys several fold over their untargeted counterparts (P=7.8×10^-5^ and 0.0020, respectively). The data herein suggest that targeting of a variety of antifungal drugs to fungal glycans may achieve lower safer effective doses and improve drug efficacy against a variety of invasive fungal infections.

## Introduction

Invasive candidiasis is among top 4 most life-threatening fungal diseases (1–5). Most *Candida species* that cause disseminated candidiasis such as *C. albicans* and *C. glabrata* are commensals that are commonly found in the gastrointestinal and urinary tracts and rarely cause invasive infections in healthy people. However immunocompromised individuals such as patients on immunosuppressants as part of cancer treatment or cell or organ transplant therapy are particularly susceptible (6–8). Candidiasis is the most common invasive fungal disease of HIV patients who developed AIDS (9, 10). Even with antifungal drug therapy, the one-year mortality rate with disseminated candidiasis ranges from 25% to 40%, depending upon the patient’s underlying conditions (3, 5, 9, 11–14). When Candida infections spread to the central nervous system and brain, the mortality rate approaches 90% (15). The annual medical costs from disseminated *Candida spp*. infections in the U.S. were recently estimated at 3 billion dollars, a third of the cost to treat all fungal diseases, and representing 45% of the U.S. hospitalizations from fungal infections (4, 16). Per patient treatment costs for candidiasis range from 40,000 to 150,000 U.S. dollars (3, 5, 16–18). Clearly, there is a considerable need for improved antifungal drug performance.

Recommended antifungals drugs to treat invasive candidiasis include the polyenes (e.g., amphotericin B, AmB), echinocandins (e.g. anidulafungin, AFG), azoles (e.g., fluconazole, fluoropyrimidines (e.g., flucytosine) or combinations of these (19–22). Most are fungicidal, while most azoles are fungistatic and genetic resistance to all but AmB are serious emerging problems. AmB was the first to be used to treat invasive candidiasis, but at effective doses and with extended treatment times, AmB and other polyenes cause renal toxicity in many patients (23–25). Because of its nephrotoxicity, AmB has been replaced by echinocandins such as AFG as the first line clinical treatment (19, 22). Lowering the effective doses of AmB and AFG, would dramatically expand our treatment options for candidiasis.

AmB is amphiphobic and quite insoluble in aqueous solutions, therefore clinical formulations often include AmB loaded into the non-polar interior of detergent micelles (e.g., AmB-DOC) or intercalated into the bilipid membrane of liposomes (e.g., un-pegylated AmBisome or our pegylated version AmB-LLs (26, 27)). Current antifungal preparations used in the clinic have the disadvantage that they deliver drug to fungal and host cells alike, and have little specificity for fungal cells. We define DectiSomes as liposomes coated with a protein that targets them to a pathogenic cell, thereby increasing drug concentrations in the vicinity of the pathogen and away from host cells (28). We previously made two classes of DectiSomes, DEC1-AmB-LLs and DEC2-AmB-LLs, by coating AmB-LLs with the carbohydrate recognition domains of Dectin-1 (27) or Dectin-2 (26). Dectin-1 (*CLEC7A*) and Dectin-2 (*CLEC4N*) are human pathogen receptors expressed on the surface of various leukocytes that recognize fungal beta-glucan and alpha-mannan containing oligosaccharides, respectively. Both glycans, the ligands for targeting by these two classes of DectiSomes are expressed in cell walls, glycoproteins, exopolysaccharide matrices, and/or biofilms of most pathogenic fungi, including *Candida spp.* (29). In vitro studies show that relative to untargeted AmB-LLs, DEC2-AmB-LLs bind to different developmental stages of *C. ablicans,* bind 100-fold more strongly, and bind primarily to oligomannans in their extracellular matrix. Furthermore, DEC2-AmB-LLs kill or inhibit the growth of *Candida* cells one to two order(s) of magnitude more effectively than AmB-LLs (26). Using a neutropenic mouse model of pulmonary aspergillosis DEC2-AmB-LLs were significanlty more effective at reducing fungal burden of *Aspergillus fumigatus* in the lungs and improving mouse survival than AmB-LLs (30). Herein, we examine the efficacy of these same DectiSomes to control *C. albicans* in a neutropenic mouse model of diseminated candidiasis. Also included are the preparations and initial examinations of AFG loaded DectiSomes and Dectin-targeted AmBisome.

## RESULTS

### DectiSomes bind efficiently to in vitro grown *C. albicans hyphae*

The binding of DEC1-AmB-LLs to *C. albicans* has not been studied in as much detail (27) as DEC2-AmB-LL binding (26). The binding of rhodamine A tagged DEC1-AmB-LLs and DEC2-AmB-LLs to *C. albicans* hyphae grown in vitro is compared in Fig. 1. By measuring the area of red liposome fluorescence from large numbers of epifluorescence images we quantified the binding data for each liposomal type. Both Dectin-1- and Dectin-2-targeted liposomes bound at least 100-fold more efficiently than our pegylated analog of AmBisome, AmB-LLs (Fig. 1D, P = 4.0×10^-6^ and 4.5×10^-6^, respectively). Bovine Serum Albumin coated liposomes, BSA-AmB-LLs, also did not bind at significant levels. The binding efficiency of the two different Dectin targeted liposomes was not statistically distinguishable (p = 0.18). However, their binding patterns differed. DEC1-AmB-LLs appeared to target exopolysaccharide distally associated with hyphae (Fig. 1A), while DEC2-AmB-LLs appeared to bind exopolysaccharide more proximally associated with hyphae and more evenly distributed throughout colonies of filamentous cells (Fig. 1B).

**Fig. 1.**
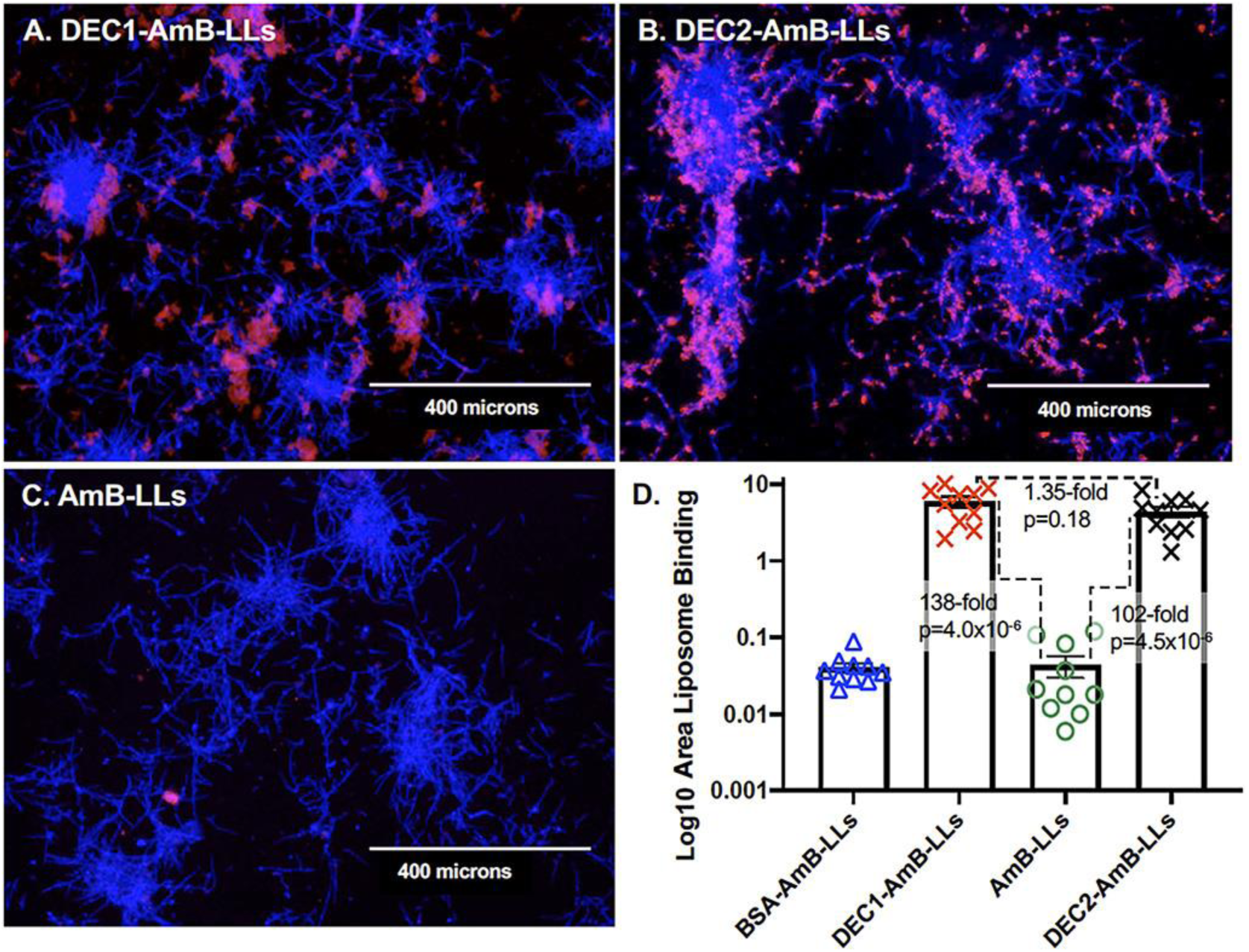
DEC1-AmB-LLs and DEC2-AmB-LLs bound specifically to in vitro grown *C. albicans* hyphae. *C. albicans* hyphae were stained with rhodamine tagged liposomes (**A**) DEC1-AmB-LLs and (**B**) DEC2-AmB-LLs, and (**C**) AmB-LLs. Binding by BSA coated BSA-AmB-LLs is not shown. **A to C**. Fungal cell chitin was stained with CW. The epifluorescence of chitin (blue) and liposomes (red) was photographed at 10X magnification. The Dectin and BSA protein concentration was 1 ug/100 uL PBS (1:100 w/v), and AmB-LLs were diluted equivalently. Size bars represent 400 microns. **D.** A scatter bar plot compares the area of red fluorescent staining quantified from 10 images for each type of liposome. Standard errors and the fold differences in average area of staining and P values are indicated for comparisons of DectiSomes to AmB-LLs.

### A neutropenic mouse model of disseminated candidiasis

We employed a neutropenic mouse model of immunosuppression to insure reproducible and sustainable invasive *C. albicans* infections were established in all mice (31–34). Neutropenic mice were infected by the intravenous injection of *C. albicans* yeast cells on Day zero (D0) and subsequently treated with an intravenous injection(s) of DEC1-AmB-LLs or DEC2-AmB-LLs, AmB-LLs, or liposome dilution buffer at various times post infection (PI). The regimens for immunosuppression, infection, treatment and assays are diagrammed in Supplemental Fig. S1. The relative effects of treating with targeted and untargeted liposomes or buffer were quantified by measuring the association of liposomes with *C. albicans* infection centers in kidneys, the fungal burden in kidneys, and mouse survival.

### DectiSomes associate with *C. albicans* infection centers in the kidneys

The goal of this experiment was to demonstrate that DEC1-AmB-LLs and DEC2-AmB-LLs are preferentially associated with *C. albicans* cells in infected kidneys as compared to untargeted AmB-LLs. Neutropenic mice were intravenously infected with 7.5×10^6^ *C. albicans* yeast cells on D0 and then given two subsequent intravenous doses of rhodamine B tagged targeted DEC1-AmB-LLs, DEC2-AmB-LLs or untargeted AmB-LLs delivering 0.4 mg/kg AmB on 3 hr PI and D1 24 hr PI (Supplemental Fig. SF1A). This amounted to 0.83 mg/kg Dectin protein per treatment for each class of DectiSomes. On D3 72 hr PI kidneys were harvested and fresh tissue was hand sectioned. Fungal chitin was stained with calcofluor white (CW) to identify infection centers and the surface of the tissue was examined top down by epifluorescence. The majority of kidney sections contained a few to a dozen CW-stained infection centers of approximately 100 to 400 microns in diameter (Fig. 2). The rhodamine red fluorescence of DEC1-AmB-LLs, DEC2-AmB-LLs and AmB-LLs, was detected in association with *C. albicans* hyphae in approximately 20%, 80%, and 5% of the infection centers, respectively (Fig. 2A, 2B, 2C), albeit, the amounts of AmB-LLs observed were often at the limit of our detection. We quantified the red fluorescent area of liposome binding within and surrounding infection centers, in images wherein liposomes were detected. A scatter bar plot (Fig. 2D) shows that respectively, DEC1-AmB-LLs and DEC2-AmB-LLs were 24-fold (P=0.027) and 56-fold (P=0.00015) more strongly associated with infection centers than AmB-LLs. This analysis gives only a semiquantitative assessment of binding, because it does not account for the differing frequency of finding infection centers with the three different types of liposomes. Replicate images of liposome binding are shown in Supplemental Fig. SF2.

**Fig 2.**
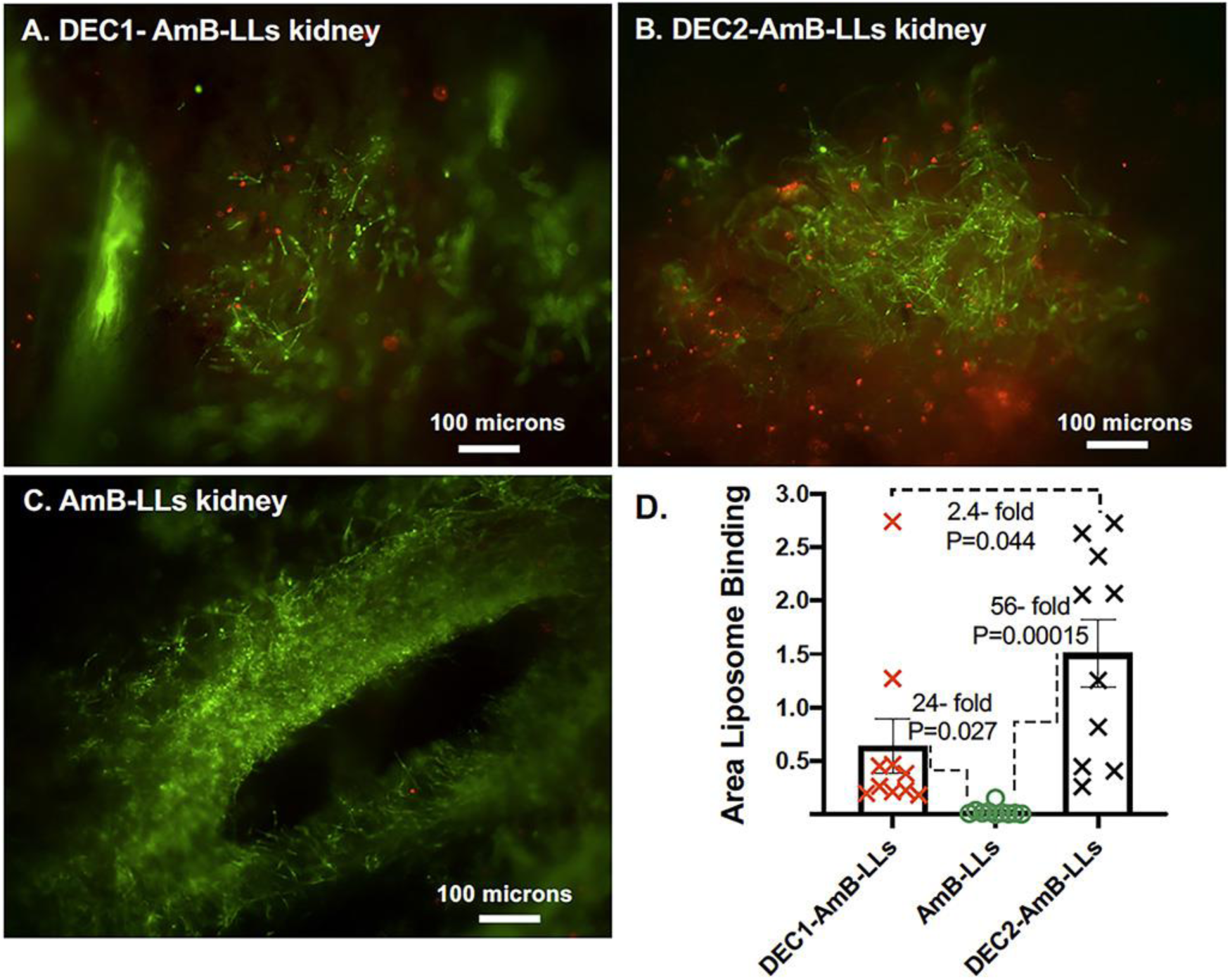
DectiSomes delivered intravenously are concentrated in *C. albicans* infection centers in the mouse kidney. Immunosuppressed mice with invasive candidiasis were injected intravenously with red fluorescent liposomes. Thick sections of the kidneys were stained with CW. **A, B. C.** The blue fluorescence of chitin (shown in green) and the red fluorescence of rhodamine tagged liposomes were photographed by epifluorescence from the surface of the tissue sections at 10X magnification. **A.** AmB-LLs. **B.** DEC1-AmB-LLs. **C.** DEC2-AmB-LLs. **D.** A scatter bar plot compares the area of red liposome fluorescence quantified from 10 images of infection centers for each treatment. Fold differences in the average area of liposome staining and P values are indicated.

### DectiSomes targeting of AmB enhanced the reduction of fungal burden in kidneys

Neutropenic mice infected with 7.5 x10^6^ *C. albicans* yeast cells and treated once 3 hr PI with either AmB-LLs, DEC1-AmB-LLs or DEC2-AmB-LLs delivering 0.2 mg/kg AmB diluted into phosphate buffered saline (PBS) or with the same amount of PBS alone (Buffer control). On D1, 24 hr PI, the mice were sacrificed and their kidneys excised, homogenized, and assayed for fungal burden. In various previous reports on neutropenic mouse models of candidiasis infected with 10^6^ *C. albicans* or 10^7^ *C. glabrata* cells, a single dose of 1.0 to 20 mg /kg AmB delivered intravenously a few hours PI as micellar AmB-DOC or liposomal AmB preparations produced 3- to 10,000-fold reductions in the kidney fungal burden relative to control mice (33–35). In our mouse model AmB-LLs delivering 0.2 mg/kg AmB provided only marginal often insignificant reductions in fungal burden relative to PBS treated mice (P=0.035 to 0.44, Fig. 3). However, mice treated with DEC1-AmB-LLs delivering 0.2 mg/kg AmB showed a 4.5-fold reduction in colony forming units (CFUs) relative to AmB-LL treated mice (P=0.013, Fig. 3A). Assays of the Relative Quantity of *C. albicans* rDNA ITS gene copies on DNA prepared from parallel samples of homogenized kidney tissue from the same mice (Fig. 3B) revealed DEC1-AmB-LL treated mice had a 6.2-fold greater reduction in fungal burden in the lungs than AmB-LL treated mice (P=4.2×10^-5^), supporting the CFU results.

**Fig. 3.**
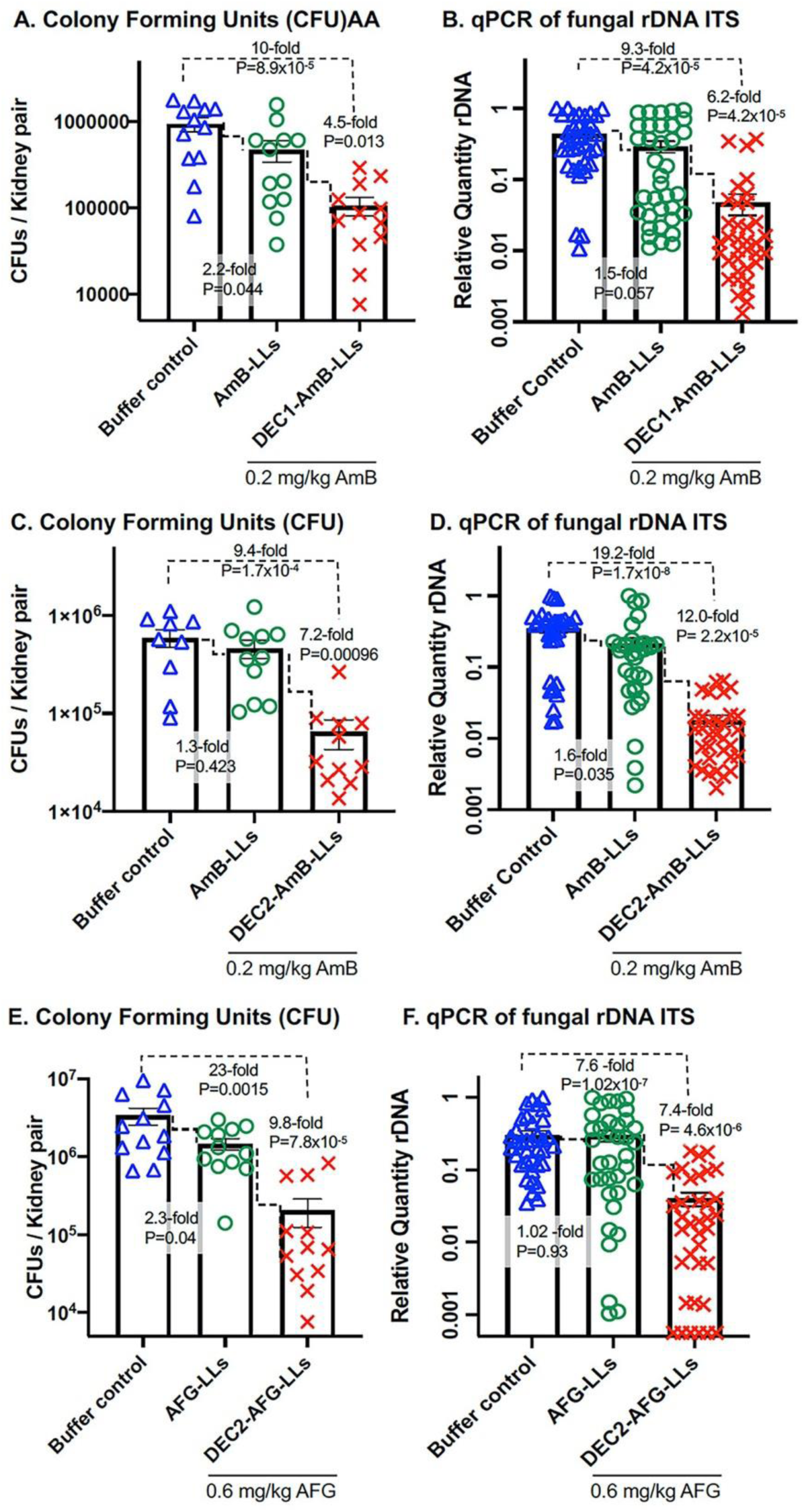
DectiSomes were more effective at reducing the burden of *C. albicans* in the kidneys as compared to untargeted AmB-LLs. Neutropenic mice infected with *C. albicans* were treated with PBS (Buffer control) or various AmB loaded liposomes. Scatter bar plots compare the fungal burden in the kidneys following treatment. **A. & B.** DEC1-AmB-LLs or AmB-LLs delivering 0.2 mg AmB/kg or PBS. **C. & D.** DEC2-AmB-LLs or AmB-LLs delivering 0.2 mg AmB/kg or PBS. **E. & F**. The performance of targeted Anidulafungin loaded liposomes. Fungal burden was assayed in infected mice treated with DEC2-AFG-LLs or AFG-LLs delivering 0.6 mg AFG/kg or PBS. CFUs or the Relative Quantity (RQ) of *C. albicans* rDNA was determined in kidney homogenates from the same mice. Twelve mice were included in each treatment group. See treatment regimens displayed in Supplemental Fig. S1A.

Mice treated with DEC2-AmB-LLs delivering 0.2 mg/kg AmB showed a 7.2-fold reduction in the kidney fungal burden relative to AmB-LLs based on CFUs (P=9.6×10^-4^, Fig. 3C) and a 12-fold reduction based on qPCR amplified rDNA ITS (P=2.2×10^-5^, Fig. 3D).

### DectiSomes targeting AFG reduced the fungal burden

AFG is a first line antifungal used to treat candidiasis, with daily patient doses of 1 to 4 mg/kg continued for several weeks (36, 37). We wished to determine if Dectin targeting might improve the performance of AFG and employed Dectin-2, to test this idea. In published studies using neutropenic mouse models of candidiasis a dose of 4 to 25 mg/kg AFG produces at least a 10-fold drop in fungal burden of in the kidneys within 24 to 48 hr (32, 38). A recent study prepared AFG-LLs loaded with 5.2 moles percent AFG relative to moles of liposomal lipid (39). In their wax moth model of candidiasis, the prophylactic administration of AFG-LLs delivering 2.6 mg/kg AFG significantly improved insect survival relative to an equivalent prophylactic dose of free AFG (39). We prepared AFG-LLs with 6.2 moles percent AFG (Supplemental Table ST1) and coated some with Dectin-2 to make DEC2-AFG-LLs. Neutropenic mice were infected intravenously with 7.5×10^6^ cells and at 3 hr PI given an intravenous dose of DEC2-AFG-LLs or AFG-LLs delivering 0.6 mg/kg AFG or liposome dilution buffer. Fungal burden in the kidneys was assayed 24 hr PI. Based on CFUs and qPCR, respectively, mice treated DEC2-AFG-LLs had a significant 9.8-fold (P= 7.8×10^-5^) and 7.4-fold (P=4.6×10^-6^) reduction in kidney fungal burden relative to those treated with untargeted AFG-LLs (Fig. 3E, 3F).

### The performance of AmBisome relative AmB-LLs

We wished to compare the performance of un-pegylated commercial AmBisome to our pegylated AmB-LLs and also to determine to what extent Dectin targeting improved the performance of AmBisome. In the neutropenic mouse model of candidiasis, we found that AmB-LLs delivering 2 mg/kg AmB reduced fungal burden in the kidneys 6.5-fold more than AmBisome delivering the same amount of AmB (P = 4.2×10^-6^, Fig. 4A). We prepared Dectin-1 and Dectin-2 coated AmBisome. Dectin-1 and Dectin-2 targeted AmBisome bound to *C. albicans* hyphae with similar specificity (Fig. 4C-4E) and efficiency (Fig. 4F) as Dectin targeted AmB-LLs (Fig. 1). When delivering 0.2 mg/kg AmB, DEC2-AmBisome reduced the kidney fungal burden 6.1-fold more than untargeted AmBisome (P=0.0125, Fig. 4B).

**Fig. 4.**
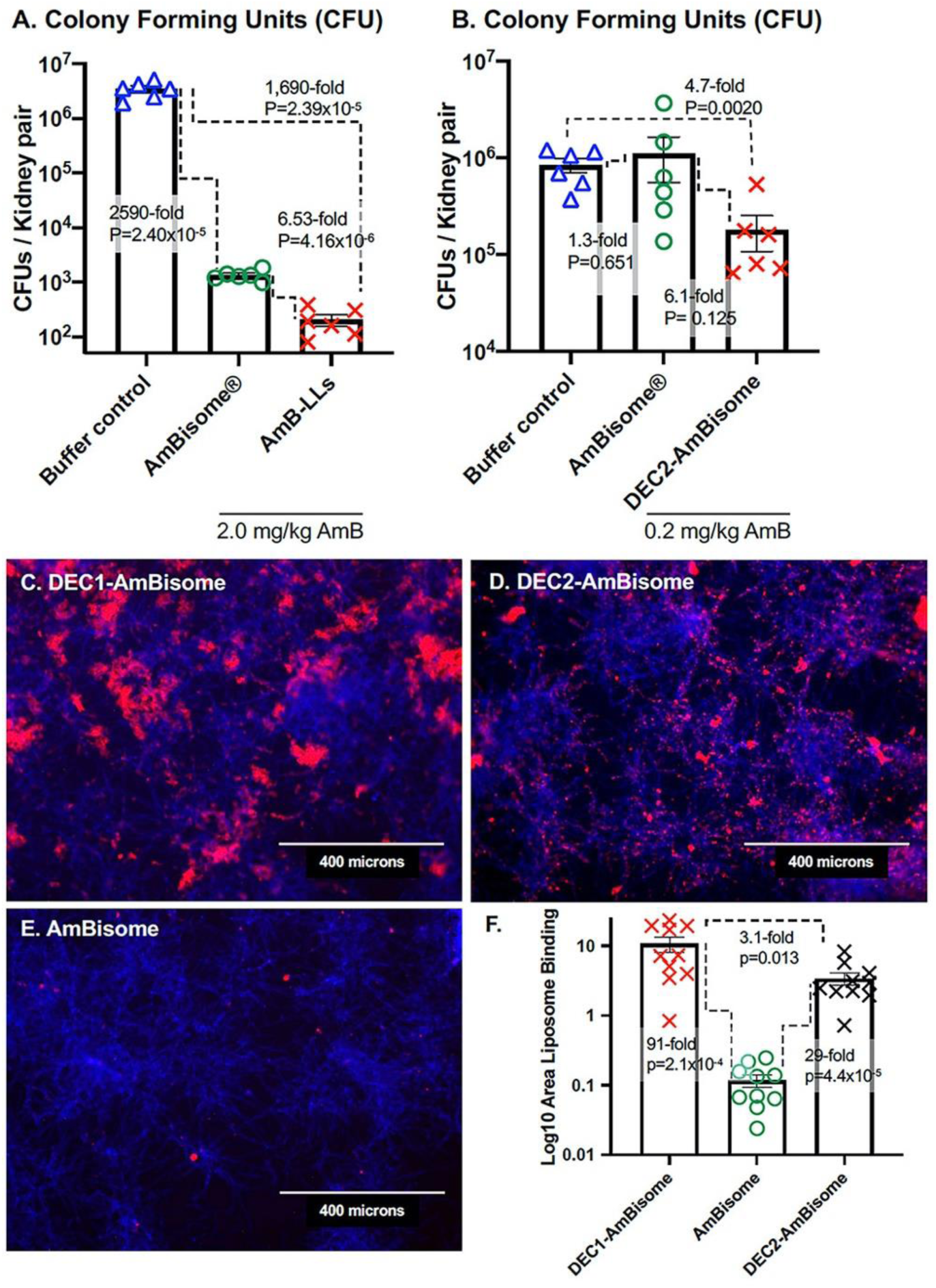
The relative performance of AmBisome. **A & B.** The kidney fungal burden was examined after neutropenic mice infected with *C. albicans* were treated with liposomes. **A.** Mice were treated with AmBisome or the AmB-LLs delivering 2.0 mg AmB/kg or PBS. **B.** Mice were treated with AmBisome and DEC2-AmBisome delivering 0.2 mg AmB/kg or PBS. See mouse treatment regimens displayed in Supplemental Fig. S1A. **C, D, E**. Fluorescent images showing the binding of Dectin targeted and untargeted rhodamine B tagged AmBisome to in vitro grown *C. albicans.* Fungal chitin was stained with CW. **F.** Quantification of the liposome binding was estimated from multiple images such as those in C-E. Standard errors, fold differences in the average area of liposome staining, and P values are indicated.

### DectiSomes increased mouse survival

Neutropenic mice were given an intravenous inoculum of 0.5 × 10^6^ *C. albicans* yeast cells, three intravenous treatments with DEC1-AmB-LLs or DEC2-AmB-LL or AmB-LLs delivering 0.2 mg AmB/kg or buffer (Control), 3 hr PI (D0), 24 hr PI (D1), and 48 hr PI (D2) (Supplemental Fig. S1B). Survival was monitored for 10 days (D10) PI as shown in Fig. 5 (31, 40, 41). All buffer-treated control mice and a few of the liposome treated mice showed reduced grooming by D3 PI. Fig. 5A presents a survival curve comparing DEC1-AmB-LLs to AmB-LLs. Forty two percent of the DEC1-AmB-LL treated mice survived to D10 as compared to 16.6% of the AmB-LL treated mice, a 2.5-fold difference in the % survival. Control mice had an average survival time of 4.6 days. DEC1-AmB-LL treatment increased the average survival time to 8.0 days as compared to 5.7 days for AmB-LL treated mice (P=0.035). Fig. 5B examines the survival of mice treated with DEC2-AmB-LLs. Sixty six percent of the DEC2-AmB-LL mice survived to D10 as compared to 8.3% of the AmB-LL mice, an 8.3-fold difference in the % survival. Control mice had an average survival time of 4.2 days, AmB-LL mice 5.6 days, and DEC2-AmB-LL mice 8.7 days, based on estimating survival time to D10. DEC2-AmB-LL treatment significantly increased the average days of survival relative to AmB-LL treatment (P=0.0006). In summary, when mice with invasive candidiasis are treated with Dectin-1 or Dectin-2 targeted DectiSomes delivering 0.2 mg/kg AmB they both showed significantly improved mouse survival relative to AmB-LL treatment. Dectin-2 appeared superior to Dectin-1 in targeting liposomal AmB.

**Fig. 5.**
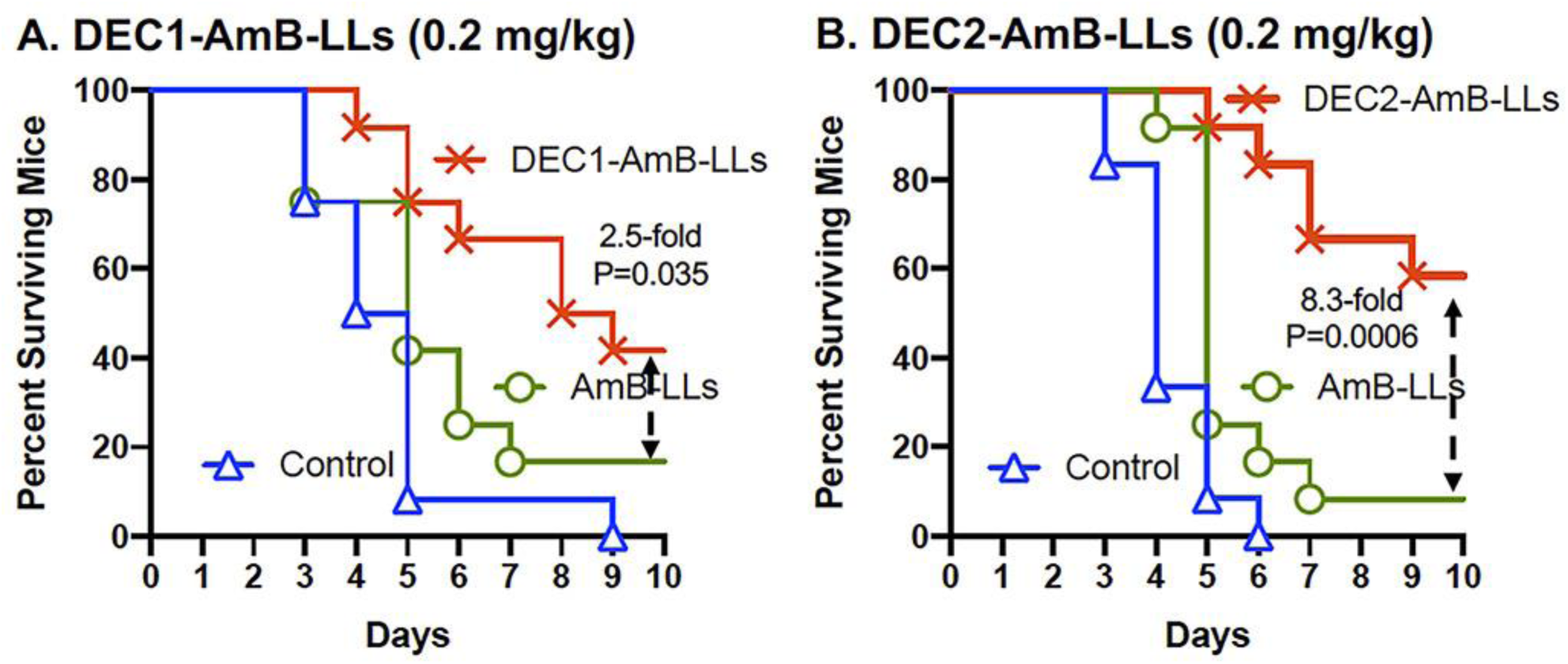
DectiSomes improved mouse survival relative to untargeted AmB-LLs. Neutropenic mice infected with *C. albicans* were treated with DEC1-AmB-LLs (**A**), DEC2-AmB-LLs (**B**) and AmB-LLs delivering 0.2 mg AmB/kg diluted into PBS or with PBS (Control). Mouse survival was monitored for 10 days PI. Fold differences in the percent of surviving mice and P values comparing the days of survival 10 days PI are indicated for the critical comparison of DectiSomes to AmB-LLs. Twelve mice were included in each treatment group. See treatment regimen displayed in Supplemental Fig. S1B.

## DISCUSSION

We showed that the AmB-LLs employed herein out performed commercial AmBisome at reducing the burden of *C. albicans* cells in the kidneys. Our AmB-LL are pegylated stealth liposomes. Pegylation protects liposomes from opsonization and phagocytosis, which significantly extends the half-life of packaged drug (42–45). Five moles percent of the liposomal lipids in the membrane of AmB-LLs are the lipid, DSPE (distearoyl-sn-glycero-3-phosphoethanolamine) coupled to poly-ethylene glycol (PEG). DSPE inserts in the liposome membrane and presents PEG on the liposome surface (27). AmBisome was first patented and introduced to the clinic before the benefits pegylation were described. Because of their pegylation, we anticipated that AmB-LLs might out-perform AmBisome. However, while they share 11 moles percent AmB, AmB-LLs and AmBisome also have different ratios of different anionic lipids and cholesterol. Hence, because of these compositional differences, we cannot conclude unambiguously that pegylation is the reason that we observed superior antifungal activity of AmB-LLs over AmBisome. Yet, our results support this concept.

DEC1-AmB-LLs and DEC2-AmB-LLs both bound to the exopolysaccharide matrix associated with in vitro grown *C. albicans* hyphae and their binding efficiency was indistinguishable. Yet, qualitatively the location of their binding in the matrix was distinct, with DEC2-AmB-LLs binding to exopolysaccharide that was more closely and uniformly associated with hyphae than DEC1-AmB-LLs. DectiSomes delivered intravenously efficiently penetrated into the kidneys of infected mice. Modest amounts of DEC1-AmB-LLs and DEC2-AmB-LLs were observed in association with infection centers, while AmB-LLs were barely detected. These data suggest that once they are bound to their glycan ligands, DectiSomes remained in place, but being unbound AmB-LLs must have been flushed out of the kidneys. However, there was a wide variation in the amount of both DectiSomes measured in individual infection centers as revealed by the wide spread of data in scatter bar plots. DectiSomes may not penetrate equivalently into all parts of the infected kidney or the glycan ligands for binding may not be equivalently expressed in all infection centers. Both types of DectiSomes appeared to be associated with hyphae, but not bound directly to them, consistent with their binding to oligoglucans and oligomannans in the associated fungal exopolysaccharide. DEC2-AmB-LLs bound significantly more efficiently to infection centers than DEC1-AmB-LLs. These results suggested the possibility that Dectin-2 targeted AmB-loaded liposomes would likely out-perform Dectin-1 targeted liposomes when their antifungal activity was tested in this mouse model of candidiasis.

Dectin-1 and Dectin-2 targeted DectiSomes delivering 0.2 mg/kg provided approximately a 9- to 19-fold reductions in kidney fungal burden relative to the buffer control and a 5- to 12-fold reduction relative to AmB-LL treated mice. DEC2-AmB-LLs appeared to be slightly superior to DEC1-AmB-LLs at reducing fungal burden relative to AmB-LLs. In mouse survival studies both Dectin-1 and Dectin-2 targeted DectiSomes delivering 0.2 mg/kg AmB improved mouse survival relative to untargeted AmB-LLs. But again DEC2-AmB-LLs appeared superior to DEC1-AmB-LLs, wherein DEC2-AmB-LLs showed relative increases the average days of survival and the percent of surviving mice.

Dectin-2 targeting of liposomal AFG (i.e., DEC2-AFG-LLs) provided a several fold increase in antifungal efficacy over untargeted AFG-LLs at reducing fungal burden. Targeting allowed a single low dose of AFG, 0.6 mg/kg, to be highly effective, a dose at which untargeted liposomal AFG-LLs was not effective. Although both AmB and AFG are amphiphobic allowing them to be intercalated into liposomal membranes, the two drugs are distinct in structure and antifungal mechanism. The polyene AmB damages the fungal membrane and osmotic integrity, while the echinocandin AFG inhibits beta-glucan synthase and ultimately oligoglucan synthesis in the cell wall and exopolysaccharide matrix. AmB is fungicidal and AFG is fungistatic. Showing improved efficacy for targeted AFG is important step forward, because it begins to generalize DectiSome targeting strategies, paving the way to improve the performance of a wide variety of other existing polyene and echinocandin drugs, other classes of antifungal drugs such as the azoles and antimetabolites, and yet to be clinically approved new drugs. Furthermore, Dectin-2 targeting improved the effectiveness AmBisome, suggesting that DectiSome targeting should improve the antimicrobial activity of nanoparticles with diverse chemical compositions, not just AmB-LLs.

Dectin-1 and Dectin-2 are both C-type lectin pathogen receptors that respond to infections by *Candida spp.* and signal the immune system of an ongoing infection. Dectin-1 is expressed primarily by neutrophils, macrophages and dendritic cells, while Dectin-2 is primarily expressed by dendritic cells. Dectin-2 appears to be the primary receptor by which Bone Marrow-derived Dendritic Cells (BMDC) signal an oligomannan dependent innate immune response to *C. albicans* yeast cells (46). BMDCs from *Clec4n^-^/Clec4n^-^* (Dectin-2 KO) mice show a hundred-fold reduction in the induction of inflammatory cytokines, such as IL-6 or TNF-alpha, when exposed to *C. albicans* cell*-*derived mannans, as compared to WT BMDCs (46). The importance and role of Dectin-1 in the response to exposure to *Candida spp.* is less clear, and appears to be less significant. There is substantial evidence that the *Candida spp.* beta-glucan ligands are heavily masked from binding either by Dectin-1 and/or anti-beta-glucan antibodies and significantly protected from the host innate immune response (47–51). For example, when BDMCs derived from *Clec7a^-^/Clec7a^-^* mice (*Dectin-1* knockout KO) are exposed to *C. albicans* or other *Candida spp.* yeast cells, their induction of inflammatory cytokines, such as IL-6 or TNF-alpha, is only reduced by 20 to 50% relative to wild type BMDCs (52). Yet, Dectin-1 KO mice infected with *C. albicans* are significantly more likely to die than WT mice, suggesting Dectin-1 does contribute positively to preventing infection. By contrast, the survival of these Dectin-1 KO mice is not reduced, when exposed to other common pathogenic *Candida spp.* such as *C. glabrata* or *C. tropicalis*, presumably due to the masking of their oligoglucans (52).

Therefore, based on the response of these two Dectins to *Candida spp*. infections and the suggestion that oligomannans were masked, we were confident at the start of this project about the potential of Dectin-2 targeting, but doubtful about the benefits of Dectin-1 targeting. We were encouraged to proceed with in vivo testing of Dectin-1 targeted DectiSomes by our strong in vitro data and modest in vivo kidney data showing DEC1-AmB-LL bound to exopolysaccharide associated with *C. albicans*. Both the partial masking of oligoglucans in vivo and the more distal association of DEC1-AmB-LLs with hyphae may explain their slightly lower effectiveness at reducing fungal burden and improving mouse survival as compared to DEC2-AmB-LLs.

*Canada spp.* form biofilms, which sequester antifungal agents and physically block access to fungal cell surfaces, thus helping them evade the host immune system and increase antifungal drug resistance (53–55). Even immunocompetent individuals may have persistent *Candida* infections, when biofilms form on implanted medical devices (53, 56, 57). AmB-LLs are significantly more effective at killing *C. albicans* residing in biofilms than either detergent solubilized micellar AmB-DOC or micellar fluconazole (58). Liposomal AmB-LLs also penetrate more efficiently into various organs (59–61), and the fungal cell wall (62), and show reduced organ toxicity and less infusion toxicity at higher AmB doses when compared to detergent solubilized AmB (23-25, 63, 64). Because of the effectiveness of liposomal formulations at both organ and biofilm penetration, new studies on therapeutics to treat candidiasis, often include variously prepared AmB-LLs or AmBisome as a standard for comparison (18, 40, 65–67), as we have done herein. Our results suggest that in neutropenic mice, DectiSomes targeted to either the oligoglucan or oligomannan components of the *C. albicans* exopolysaccharide matrix enhance the performance of AmB-LLs. Future studies need to focus specifically on the efficacy of DectiSomes against various *C. albicans* biofilms.

### Conclusions

Dectin-1 or Dectin-2 targeting of liposomal AmB to *C. albicans* glycans significantly improved the performance of AmB, over untargeted AmB-LLs. Similarly, Dectin-2 targeting improved the performance of commercial AmBisome and of liposomal AFG. These data suggest targeting may improve the performance of a wide variety of drugs packaged in diversely structured nanoparticles. Oligoglucans and oligomannans, the respective glycan targets of Dectin-1 and Dectin-2, are ubiquitous components of the cell wall and biofilms of most pathogenic fungi suggesting there is pan-antifungal potential for DectiSome technology. A previous study showed that oropharyngeal delivery of Dectin-2 targeted DectiSomes to neutropenic mice with pulmonary aspergillosis was more effective at treating the infection than AmB-LLs (30). Herein, we show that DectiSomes delivered intravenously were effective at controlling an invasive *Candida* infection and penetrated into a host organ, the kidneys. Hence, it appears that DectiSomes have considerable potential as pan-antifungal agents whether delivered directly to the lungs or delivered intravenously. The improved performance of DectiSomes needs to be tested in a variety of other mouse models of fungal diseases, such as cryptococcal meningitis and pulmonary mucormycosis and superficial infections such as keratitis, and tested with antifungal drugs other than AmB and AFG.

## MATERIALS AND METHODS

### Strains and culture

*C. albicans* strain SKY43, expresses GFP under control of the ADH1 promoter (68) and was derived from a human isolate (SC5314, ATCC MYA-2876) deleted for URA3 (strain CA14, Δura3::imm434/Δura3::434)(69). *C. albicans* yeast cells were grown to early log phase in YPD, washed once into fresh YPD, aliquoted, snap frozen in liquid nitrogen, and stored frozen at -80°C in 25% glycerol. Cells were thawed once or twice just before use, vortexed, and diluted to the desired cell concentration in sterile saline. The viability of the thawed cultures was close to 99%. Mice were infected via the retroorbital injection of 100 uL of saline containing 7.0 × 10^6^ or 0.5 × 10^6^ yeast cells (67) (Supplemental Fig. S2).

Seven- to eight-week-old outbred female CD1 (CD-1 IGS) Swiss mice (27 g to 30 g ea.) were obtained from Charles River Labs. Mice were maintained in UGA’s Animal Care Facility. All mouse protocols met guidelines for the ethical treatment of non-human animals outlined by the U.S. Federal government (70) and UGA’s Institutional Animal Care and Use Committee (AUP #A2019 08-031-A1).

### In vitro binding studies

For in vitro binding studies 10,000 cells/mL *C. albicans* yeast cells were plated in 500 uL of RPMI 1640 media lacking red dye at pH 7.5 in each well of a 24 well microtiter plate and grown for 12 hr to achieve approximately 50% coverage with hyphae. Cells were washed once with PBS, fixed in 4% formalin for 45 min, and washed 3x with PBS. Cells were blocked with PBS + 5.0% BSA for 30 min, treated with rhodamine red fluorescent liposomes in this blocking buffer, stained with 25 uM CW (Blankophor BBH SV-2560; Bayer, Corp.) for 60 min, and washed 3x with the same buffer. Images were taken on an EVOS imaging system using the DAPI and RFP fluorescent channels and the red fluorescence area within un-enhanced images was quantified in ImageJ (26). The accompanying images presented were enhanced equivalently in the blue and red channels.

### Neutropenic model of disseminated candidiasis

Immunosuppressed neutropenic mice were obtained by treatment with both the antimetabolite cyclophosphamide (CP, Cayman #13849) and the synthetic steroid triamcinolone (TC, Millipore Sigma # T6376) following the schedules shown in Supplemental Fig. SF1. Five or six mice were in each treatment group and in some cases two replicate experiments. CP and TC stocks, dilutions, and injection methods were described recently (30).

Infected buffer control animals not receiving antifungal therapy first showed a ruffled coat due to reduced grooming, then decreased movement, followed by abnormal posture, trembling and severe lethargy. The onset of symptoms occurred much more rapidly in animals receiving the larger fungal inoculum size and was reduced in animals receiving liposomal AmB. Once mice showed severe lethargy and were moribund, they were sacrificed by cervical dislocation following anesthesia with isoflurane (Animal Use Protocol, A2019 08-031-A1).

### Liposomes and drugs

We constructed AmB-LLs, DEC1-AmB-LLs, DEC2-AmB-LLs, and BSA-AmB-LLs as described previously (26, 27). Dectisomes contain 1 mole percent dectin relative to moles of liposomal lipid. Similar to AmBisome they contain 11 moles percent AmB, but our liposomes also contain two moles percent Rhodamine B-DHPE for visualization.

AFG-LLs were prepared in 273 μL batches using a remote loading method analogous to that which we used to prepare AmB-LLs (26, 27). To quantify AFG loading into liposomes, we determined that AFG had an extinction coefficient (17.4 O.D. /mg/mL) at A340 in DMSO, using a dilution series and a Bio-Tek Synergy HT microtiter plate reader (Supplemental Fig. SF3). Ten moles percent AFG (1.7 mg, 1.5 μmoles) relative to moles of liposomal lipid was dissolved in 13 uL DMSO and added to 15 μmoles liposomal lipid (260 μL of 100 nanometer diameter Formumax liposomes (F10203, Plain)). AFG and liposomes were incubated for 72 hours at 37°C with gentle tumbling. The AFG-LLs were spun at room temperature for 2 min at 1,000 × g to sediment the remaining insoluble AFG that was not loaded into liposomes and did not remain soluble between liposomes. We had predetermined that the solubility of AFG in our liposome loading buffer (10% sucrose, 20 mM HEPES, pH 7.0-7.5, and 5% DMSO) to be 0.31 mg/273 μL). The AFG precipitate (i.e., that not loaded in liposomes) was then dissolved in DMSO and quantified at A340. By subtraction of the insoluble AFG and the predetermined solubility in loading buffer, we calculated that the AFG-LLs contained 6.2 moles percent AFG. One mole percent Dectin-2 modified with a lipid carrier, DEC2-PEG-DSPE, was then added to the AFG-LLs to make DEC2-AFG-LLs (26, 27). See Supplemental Table ST1.

### Liposome binding to infection centers in the kidneys

Hand sections of freshly harvested kidney were cut and stained for fungal chitin with calcofluor white (Blankophor BBH SV-2560; Bayer, Corp.) as previously described (30). The blue fluorescent chitin ex360/em470 and rhodamine tagged liposomes ex560/em645 were examined by epifluorescence microscopy using a LEICA DM6000 compound fluorescent microscope at 10X magnification as described previously for liposome binding to lung tissue (30). The area of red fluorescent liposome binding in the original TIFF images was quantified in ImageJ as described previously (30). For photographic presentation of binding the blue CW and red liposome channels of the original Tiff images were equivalently enhanced in Photoshop (version 20.0.8) to aid in visualizing fungal cells and liposomes, respectively, and then converted to JPEG images for presentation.

### Fungal burden estimates

Fungal burden was estimated in excised kidney pairs from infected animals on D1 PI by assaying both the number of CFUs and the amount of *C. albicans* ribosomal rDNA intergenic transcribed spacer (ITS) estimated by quantitative real time PCR (qPCR). Kidney pairs were weighed and minced into hundreds of approximately 1 mm^3^ pieces, the pieces mixed to account for the uneven distribution of infection centers, and aliquoted into 25 mg samples. *CFUs.* 25 mg of the minced kidney tissue was homogenized for 60 seconds in 200 uL of PBS using a hand-held battery powered homogenizer (Kimble, cat#749540-0000) and blue plastic pestle (Kimble Cat#749521-1500). The homogenate was spread evenly by shaking with sterile glass beads on 5 mm thick YPD (yeast extract, peptone, and dextrose) agar plates containing 100 ug/mL each of Kanamycin and Ampicillin. After a 11 hr incubation at 37°C, the microcolonies of 5 to 300 microns in diameter were counted on an EVOS imaging system (AMG Fl) at 4X magnification. An example of the images of microcolonies used to make CFU estimates is shown in Supplemental Fig. SF4. The number of CFUs was corrected for the area of the entire plate relative to each microscopic field and the weight of each kidney pair. The numbers of colonies were often so low for the DEC1-AmB-LL and DEC2-AmB-LL treated samples that 20 or more fields had to be counted on each plate to record a statistically significant number of colonies. *qPCR.* DNA was extracted from 25 mg parallel samples from kidney homogenates using Qiagen’s DNeasy® Blood & Tissue Kit (#69504) modified as we described previously for *A. fumigatus* infected lung tissue (30). We typically obtained 25 TI 40 ug of total DNA from 25 mg of kidney tissue. Quantitative real-time PCR (qPCR) was used to estimate the amount of *C. albicans rDNA* ITS sequence in 100 ng samples of infected kidney DNA using the conditions described previously (30). Several new PCR primer pairs designed to amplify the ITS downstream of 18S rDNA of *C. albicans* were designed and tested against purified *C. albicans* DNA. The optimal primer pair giving the lowest cycle threshold value (Ct) and a single dissociation peak had the following sequences (forward primer Ca18S-4S, 5’-TAGGTGAACCTGCGGAAGGATCATTA and reverse primer Ca18S-2A 5’-TTGTAAGTTTAGACCTCTGGCGGCA). This primer pair gave no detectable product even after 45 cycles of PCR, when uninfected kidney tissue DNA was examined. The Relative Quantity (RQ) of *C. albicans rDNA ITS* was determined by normalizing all Ct values to the lowest Ct value determined for infected control kidneys using the dCt method (71).

### Data management

Raw quantitative data were managed in Excel v16.16.27. Scatter bar plots, survival plots, and XY plots were prepared in GraphPad Prism v.9.0.0. Because the data for liposome binding, fungal burden estimates and days of survival were reasonably normally distributed, the Student’s two tailed t-Test was used to estimate P values (72).

## ACKNOWLEDGMENTS

We wish to thank Kristine Wilcox and the other staff members of UGA’s University Research Animal Facilities for the conscientious care of our mice.

## Supplemental Tables and Figures

**Supplemental Table ST1.** Chemical composition of Anidulafungin loaded liposomes (AFG-LLs).

| Compound | µg in each preparation of liposomes | µg / µmole Mol. Wgt. | µmoles total | Moles percent relative to moles of liposomal lipid |
| --- | --- | --- | --- | --- |
| DSPC (1,2-distearoyl-sn-glycero-3-phosphocholine) | 6516.00 | 790.00 | 8.25 | 50.00 |
| cholesterol | 1591.00 | 387.00 | 4.11 | 45.00 |
| mPEG2000-DSPE | 2302.10 | 872.00 | <u>2.64</u> | <u>5.00</u> |
| Total mmoles and moles percent liposomal lipid |  |  | 15.00 | 100.00 |
| Anidulafungin (AFG) | 1030.00 | 1140.00 | 0.90 | 6.02 |
| Lissamin Rhodamine B 1,2-Dihexadecanoyl-sn-Glycero-3-Phosphoethanolamine, Triethylammonium Salt | 400.00 | 1333.81 | 0.30 | 2.00 |

**Supplemental Fig. SF1.**
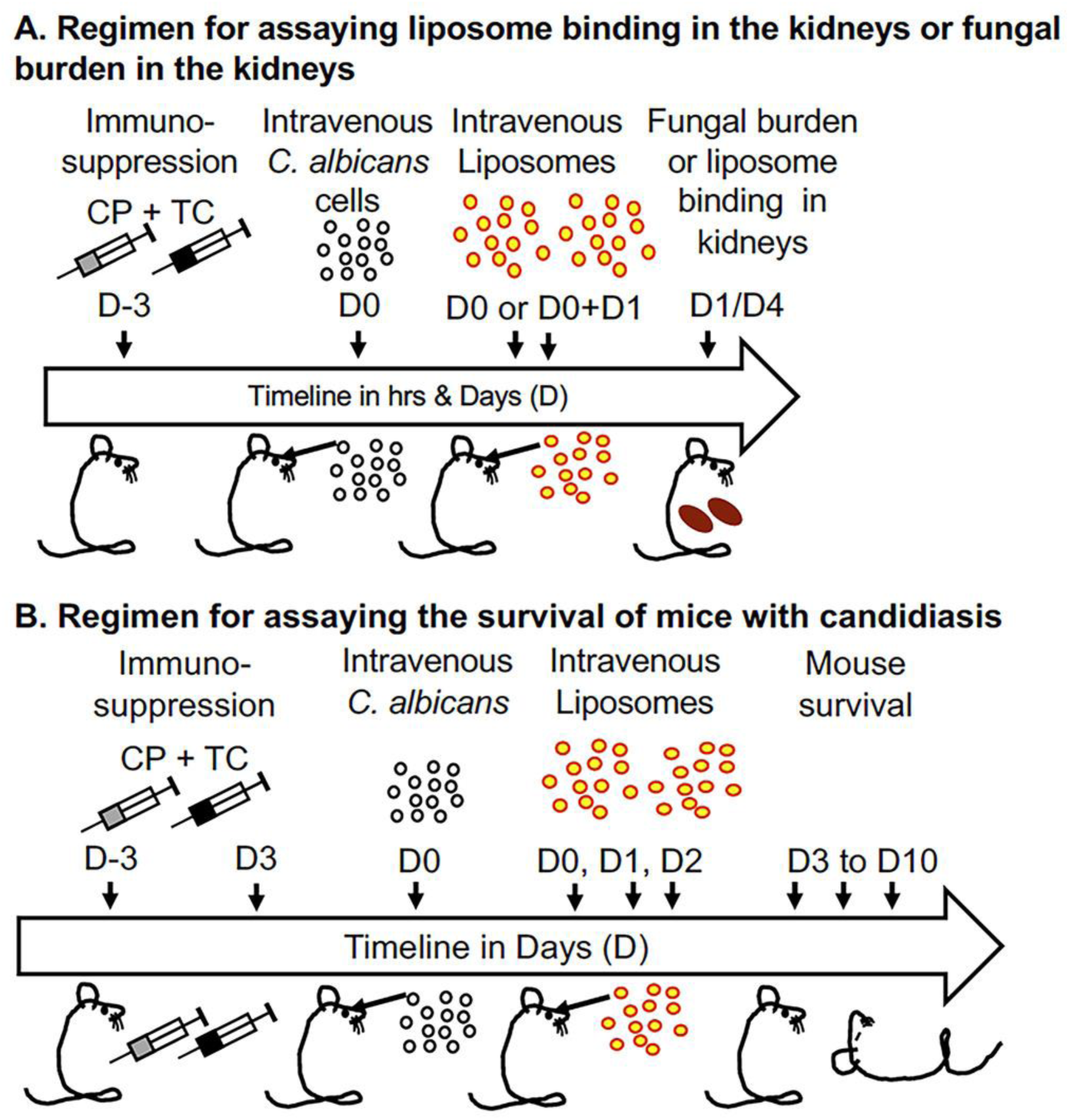
Treatment regimens used to assay liposome binding, fungal burden and mouse survival-immunosuppression, infection, liposome treatments and endpoints. A neutropenic mouse model was used to ensure reproducible infection by *C. albicans* yeast cells. **A.** The regimen used to assay binding of DectiSomes to infection centers in the kidneys as compared to AmB-LLs and the impact of DectiSomes on fungal burden as compared to untargeted liposomes or control buffer. Liposome binding was assayed on Day 4 (D4) PI and fungal burden was determined on D1 PI. **B.** Regimen used to assay mouse survival after treatment with DEC1-AmB-LLs, DEC2-AmB-LLs, AmB-LLs or control buffer. The Day(s) (D) of treatment are indicated before and after the day of infection (D0). CP, cyclophosphamide. TC, triamcinolone. Survival was monitored until D10.

**Supplemental Fig. SF2.**
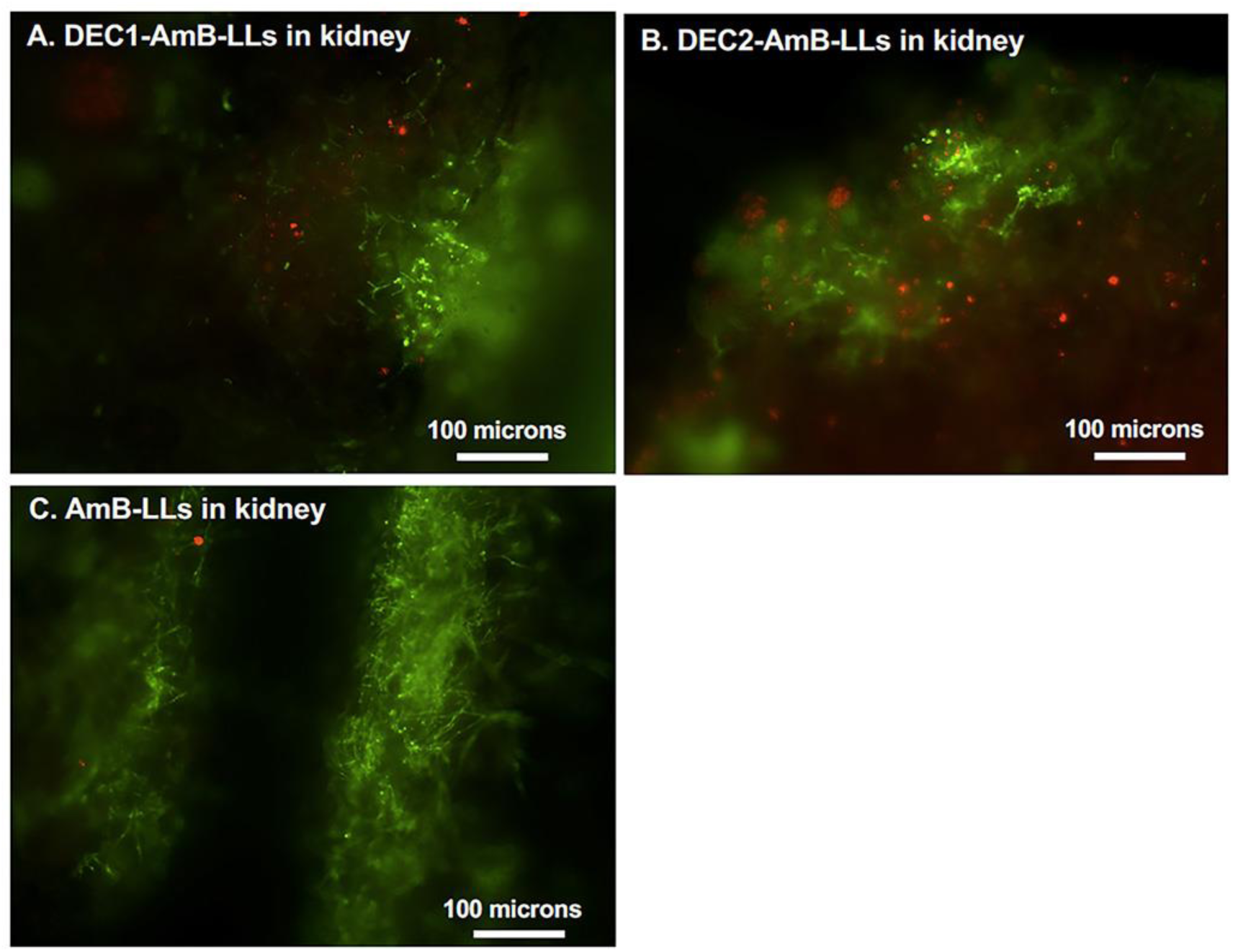
DectiSomes targeted by Dectin-1 or Dectin-2 and delivered intravenously are concentrated in *C. albicans* infection centers in the mouse kidney. Replicate images for the experiment illustrated in Fig. 2 showing the superior binding of DEC1-AmB-LLs and DEC2-AmB-LLs to *C. albicans* colonies in the kidneys relative to untargeted AmB-LLs.

**Supplemental Fig. SF3.**
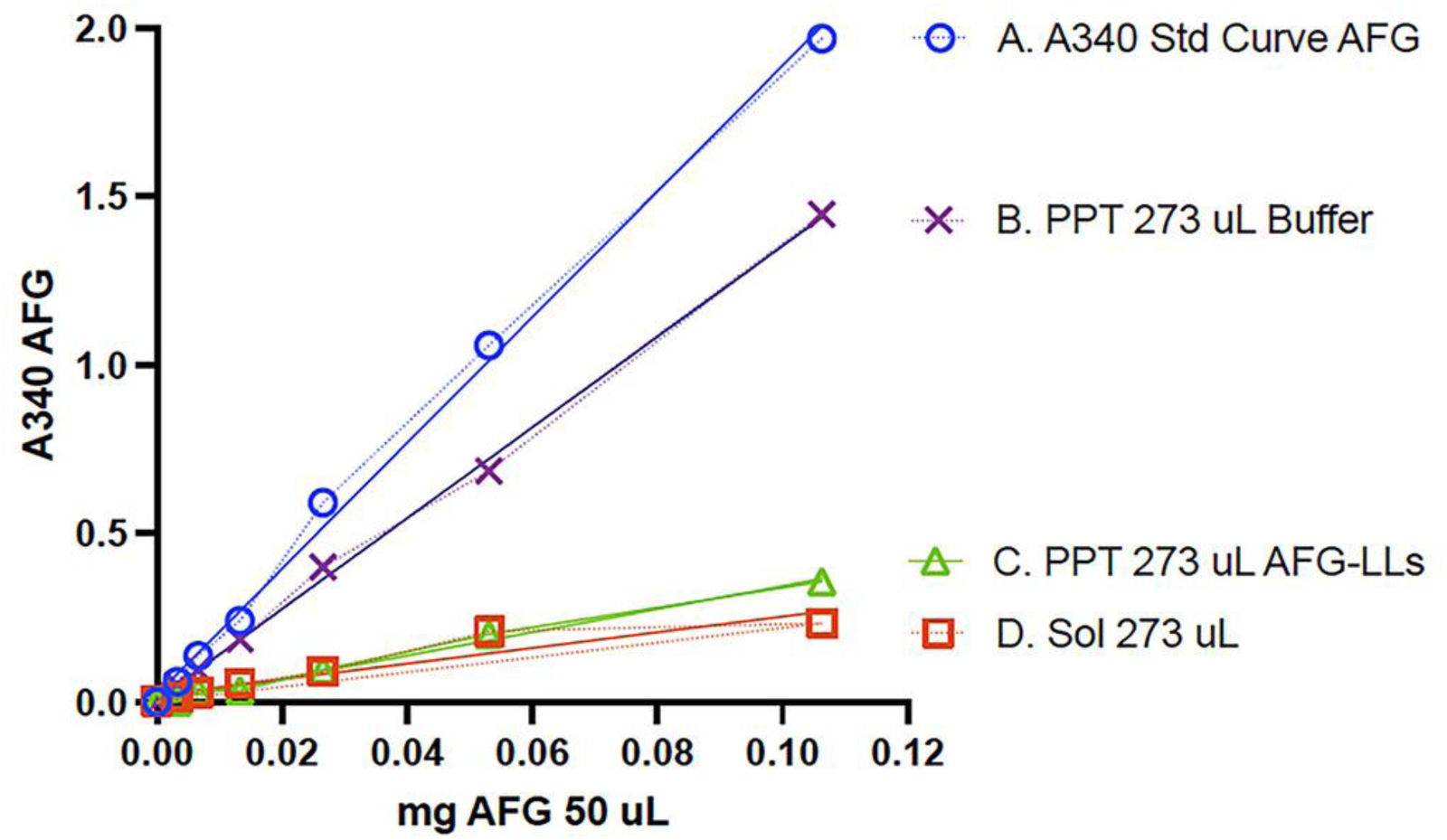
Experimental data used to quantify the moles percent of AFG loaded into AFG-LLs. Light scattering from liposomes prevents a direct measurement of the UV absorbance of drugs loaded into liposomes. Hence, we estimated the percent of AFG loaded into liposomes by the subtraction of that which was not loaded into liposomes as we have done before to estimate the loading of AmB (27). **A.** Standard curve plotting the absorbance of AFG at A340 in a dilution series in a 96 well microtiter plate vs the amount mg amount of AFG dissolved in 50 uL of DMSO. When more than 0.12 mg of AFG were examined the A340 absorbance values were too high to be read. **B.** When 1.7 mg of AFG was tumbled for 3 days in 273 uL of liposome buffer, was 81.3% of the total remained insoluble. This was determined by taking the insoluble AFG precipitate and dissolving it in 50 uL DMSO and reading A340 for a dilution series. **C.** After incubating liposomes and 1.7 mg of AFG together in 273 uL the insoluble AFG was spun down and assayed by in a dilution series in 50 uL DMSO. This is the amount of AFG not taken up by liposomes and not soluble in the buffer surrounding them. **D.** The amount of AFG that remained soluble in 273 uL of liposome buffer was estimated by subtracting the data in curve B from that in A.

**Supplemental Fig. SF4.**
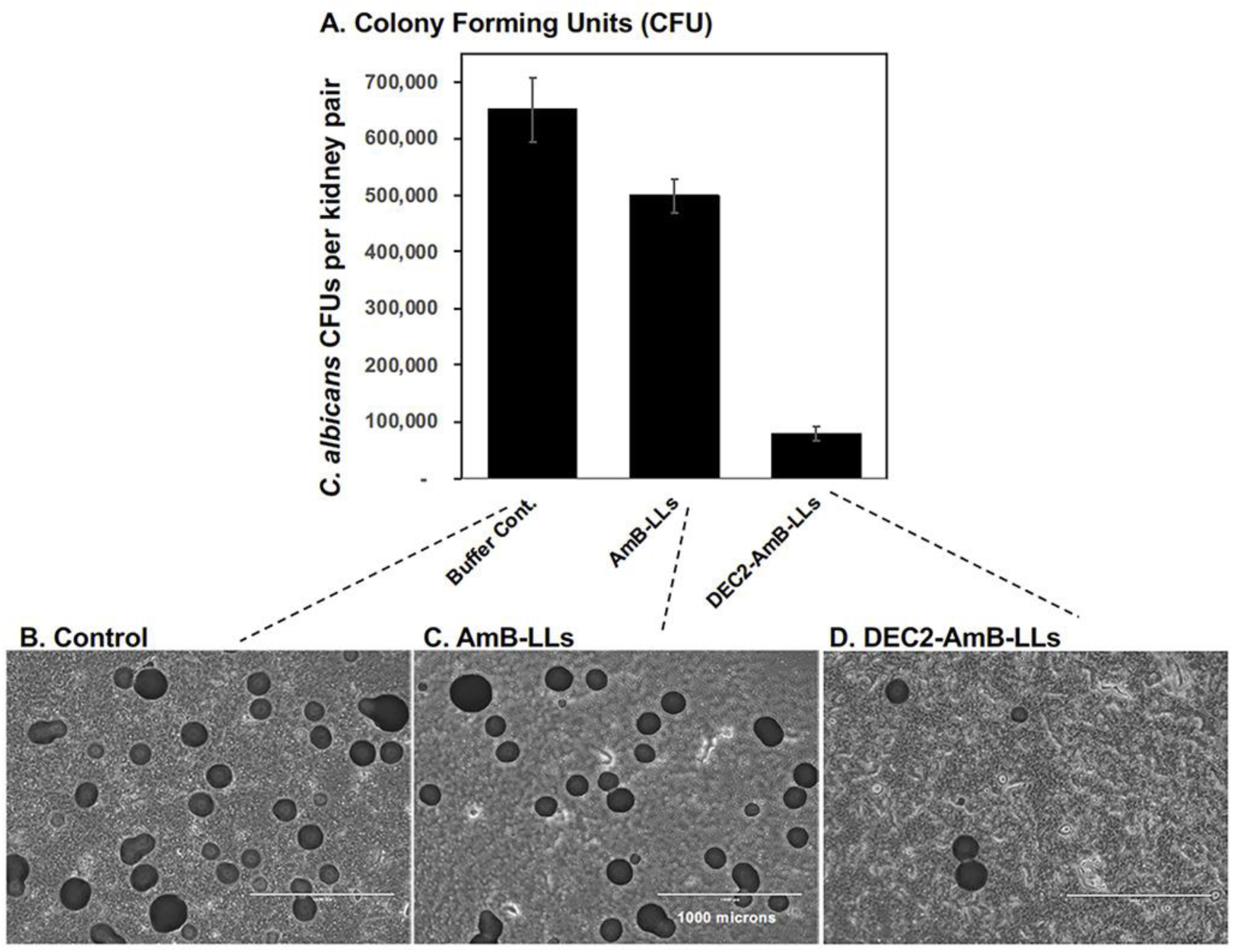
Assays of *C. albicans* microcolonies to determine fungal burden in the kidneys of neutropenic mice with candidiasis than AmB-LLs (an example experiment). Aliquots of homogenized kidney tissue were diluted into PBS, plated on YPD agar, incubated 11 hr at 37°C, and microcolonies counted. **A.** A bar plot compare the average number of CFUs of *C. albicans* for mice treated once with DEC2-AmB-LLs or AmB-LLs delivering 0.2 mg/kg AmB or with liposome dilution buffer. Standard errors are indicated by a line and whisker. **B, C, & D.** Examples of the images used to make CFU estimates. Microcolonies ranging from 5 to 300 microns in diameter were counted from the bottom of agar petri plates on an EVOS imaging system at 4X magnification. The number of CFUs was corrected for the area of the entire plate relative to each microscopic field, the amount of homogenized kidney tissue plated, and the weight of each kidney pair. Six mice were in each treatment group in this example.

